# A novel hybrid Wireless Integrated Sensing Detector for simultaneous EEG and MRI (WISDEM)

**DOI:** 10.1101/2024.07.31.606016

**Authors:** Yi Chen, Wei Qian, Daniel Razansky, Xin Yu, Chunqi Qian

## Abstract

Concurrent recording of EEG/fMRI signals reveals cross-scale neurovascular dynamics that are crucial for elucidating fundamental linkage between function and behaviors. However, MRI scanners generate tremendous artifacts for EEG detection. Despite existing denoising methods, cabled connections to EEG receivers are susceptible to environmental fluctuations inside MRI scanners, creating baseline drifts that complicate EEG signal retrieval from the noisy background. Here, a **W**ireless **I**ntegrated **S**ensing **D**etector for simultaneous **E**EG and **M**RI (WISDEM) is developed to encode fMRI and EEG signals on distinct sidebands of the detector’s oscillation carrier wave for detection by a standard MRI console over the entire duration of fMRI sequence. Local field potential (LFP) and fMRI maps are retrieved through low-pass and high-pass filtering of frequency-demodulate signals. From optogenetically-stimulated somatosensory cortex, the positive correlation between evoked LFP and fMRI signals validates strong neurovascular coupling, enabling cross-scale brain mapping with this 2-in-1 transducer as a research and diagnostic tool.

## Introduction

A major challenge in brain research is to link functional perspectives across scales, from the cellular level to the circuit and eventually to the systems level. Functional Magnetic Resonance Imaging (fMRI) has been developed to indirectly map neuronal activity across the entire brain, based on vascular hemodynamics (*e.g.*, blood flow, blood volume, or blood oxygenation levels) which contribute to fMRI signals^1–5^. To characterize the neuronal basis of the fMRI signal, the MR-compatible EEG preamplifier has been implemented inside the MRI scanner during scanning^6^, setting an ingenious scheme for concurrent acquisition of fMRI and EEG signals. Echoing this pioneering work, simultaneous electroencephalogram (EEG) and fMRI have been increasingly utilized to monitor both neuronal and hemodynamic activities^7–18^, elucidating the neurovascular coupling mechanisms underlying the hemodynamic response detected by fMRI in healthy and diseased brains^19–21^. EEG can provide temporal information about brain state dynamics while fMRI can map spatial distribution of brain state network. When used in combination, EEG/fMRI have enhanced capability to localize disease foci^22,23^ and to monitor brain network evolution during resting state^24–30^, sleep^31,32^ or cognition^33–35^.

Despite its steady progress over the past two decades, simultaneous EEG/fMRI is still technically demanding in research and clinical settings. To record tiny EEG signals inside an MR scanner, reliable cable connections are required to create common voltage ground between the skull, the EEG receiver and even the scanner bore^20^. This multi-cable configuration requires careful installation, because even slight changes in junction resistance may lead to noticeable changes in baseline, making it challenging to reproducibly retrieve a baseline pattern that is necessary for follow-up analysis. Also, the wired connections between recording electrodes and EEG receiver collect electromagnetic interference artifacts^36–38^, especially during the MR excitation pulses and switching magnetic field gradients^39–41^. Although these artifacts can partially be removed during post-processing^42–51^, reliable recovery of weak EEG signals from the much stronger background interference requires concurrent use of high-gain preamplifiers and high-speed Analog Digital Converters with large dynamic range^52^, necessitating the use of extra shielding and powering modules, making the whole system complex and with safety concerns^53–56^. An alternative way for artifact reduction is to synchronize the EEG apparatus with the MR-scanner, so that EEG signals are acquired only during the MRI acquisition window when the excitation pulse is switched off and the encoding gradient remains stable^57^. However, this approach requires precise synchronization with the MR-scanner and prior knowledge about the pulse sequence, which is difficult to implement. Recently, wireless electrophysiology transducers^58,59^ have been developed to utilize on-board gradient sensors and micro-controllers for dynamically identifying proper acquisition windows with stable magnetic field gradients, enabling the synchronically acquired EEG signals to be encoded onto the wireless carrier wave and detected by a standard MRI coil. Promising as it is, this approach requires on-board microcontrollers and dedicated RF transmitters that are powered by sizable internal batteries, making them hard to miniaturize for interventional and implantable applications. All these technical challenges preclude widespread usage of simultaneous EEG/fMRI in neuroscience research and clinical diagnosis.

To overcome above-mentioned limitations, in this study we have developed a wirelessly powered oscillator that can simultaneously encode fMRI and EEG signals. This is a major improvement over our previous work on Wireless Amplified NMR Detectors (WAND) for deep tissue imaging^60^ that combined wirelessly powered amplifiers^61^ with miniaturized micro-coils^62,63^. Here, when the wireless pumping power is increased beyond an oscillation threshold^61^, the WAND becomes an oscillator that can directly convert wirelessly provided pumping power into sustained oscillation currents near the circuit’s resonance frequencies. Unlike conventional voltage-controlled oscillators that can only encode low-frequency signals after down-conversion^64^, the wireless oscillator can utilize circuit nonlinearity to combine down conversion and frequency encoding of MRI signals into a single stage^65^. Because circuit oscillation can also be modulated by low-frequency bias voltages applied on its nonlinear components^66^, low-frequency neuronal signals are also encoded onto the same frequency-modulated (FM) carrier wave, but on a distinct sideband from the simultaneously encoded high-frequency MRI signals.

Importantly, the oscillation carrier wave can be wirelessly detected by a standard MRI coil and recorded by the MR scanner over the entire duration of MR acquisition windows, in the same way as how conventional MR signals are detected. Without the need for dedicated gradient sensors or synchronization apparatus, the oscillator can reliably encode MRI and EEG signals, even during gradient switching periods. Since the down-converted MRI signals and neuronal signals exhibit different frequency separations from the carrier center, they can be told apart by high-pass and low-pass filtering following frequency demodulation. As a result, no dedicated hardware is needed to synchronize MRI and EEG detection. The pumping power can reduce the circuit’s effective resistance and increase its quality factor by ∼39000 fold, making the circuit’s oscillation frequency very sensitive to small modulation voltages, thus obviating the need for high-power preamplifiers or digitizers that were traditionally required to recover subtle neuronal signals from the artifactual background. Without the need for ADC converters or microprocessors, the oscillator has a compact design that is easy to implant on the skull, requiring only a few milliwatts of wireless power to activate the transducer, inducing negligible heating effects.

In this study, we first fabricated a **W**ireless **I**ntegrated **S**ensing **D**etector for simultaneous **E**EG and **M**RI (WISDEM) by retrieving low and high frequency signals from FM-encoded sidebands. Then we tested the feasibility and performance of WISDEM in phantoms to retrieve low-frequency voltage signals when a train of sinusoidal waves were directly injected into the sensing electrodes. Furthermore, we validated the detector’s imaging performance by observing robust Echo Planar Imaging (EPI)-BOLD in the S1 forepaw region (S1FP) when the rodent was given electrical forepaw stimulation. Lastly, we combined optogenetic stimulation with simultaneous acquisition of local field potential and fMRI signals to expand the applicability of WISDEM. These results demonstrate the utility of this hybrid 2-in-1 detector for simultaneous EEG and fMRI signal recordings of rodent brains through the MR console system, paving the way of cross-bandwidth neuronal and hemodynamic signals.

## Results

### 1. Operation Principle and Circuit Fabrication of the WISDEM

The parametric resonator (PR) is a circular-shaped loop-gap resonator with a continuous center conductor to bridge its virtual grounds. As a result, the resonator has a <u>b</u>utterfly resonance mode (**Fig. 1a**) at a lower frequency *ω_br_* and a <u>c</u>ircular resonance mode (**Fig. 1b**) at a higher frequency *ω_cr_*. When a pumping signal is applied at approximately the sum frequency of these two modes *ω_br_*+*ω_cr_*, the resonator can oscillate at frequencies (*ω_b_* and *ω_c_*) that are close to the resonance frequencies (*ω_br_* and *ω_cr_*) of individual modes, *i.e.*, *ω_b_ ∼ ω_br_* and *ω_c_ ∼ ω_cr_*. Once the pumping signal *ω_p_* is determined by an external frequency synthesizer, it will also determine the sum of butterfly and circular oscillation frequencies, *i.e.*, *ω_p_ = ω_b_ + ω_c_*. If the butterfly mode oscillation signal falls within the detection band of the MRI scanner, it can be detected by a standard MRI coil. As explained in the Appendix, the oscillation frequency *ω_b_* has a linear relation with the resonance frequency *ω_br_*. If at the same time, the butterfly mode also interacts with an MRI signal that is separated from *ω_b_* by an offset Δ*f* that is smaller than the imaging bandwidth, the MRI signal will interact with the oscillation signal, creating a down-converted signal at Δ*f* that can frequency-modulate the oscillation signal at the same time^65^. In this way, both the low-frequency EEG signal and the high-frequency MRI signal can be encoded onto the same carrier wave for wireless transmission.

**Fig. 1.**
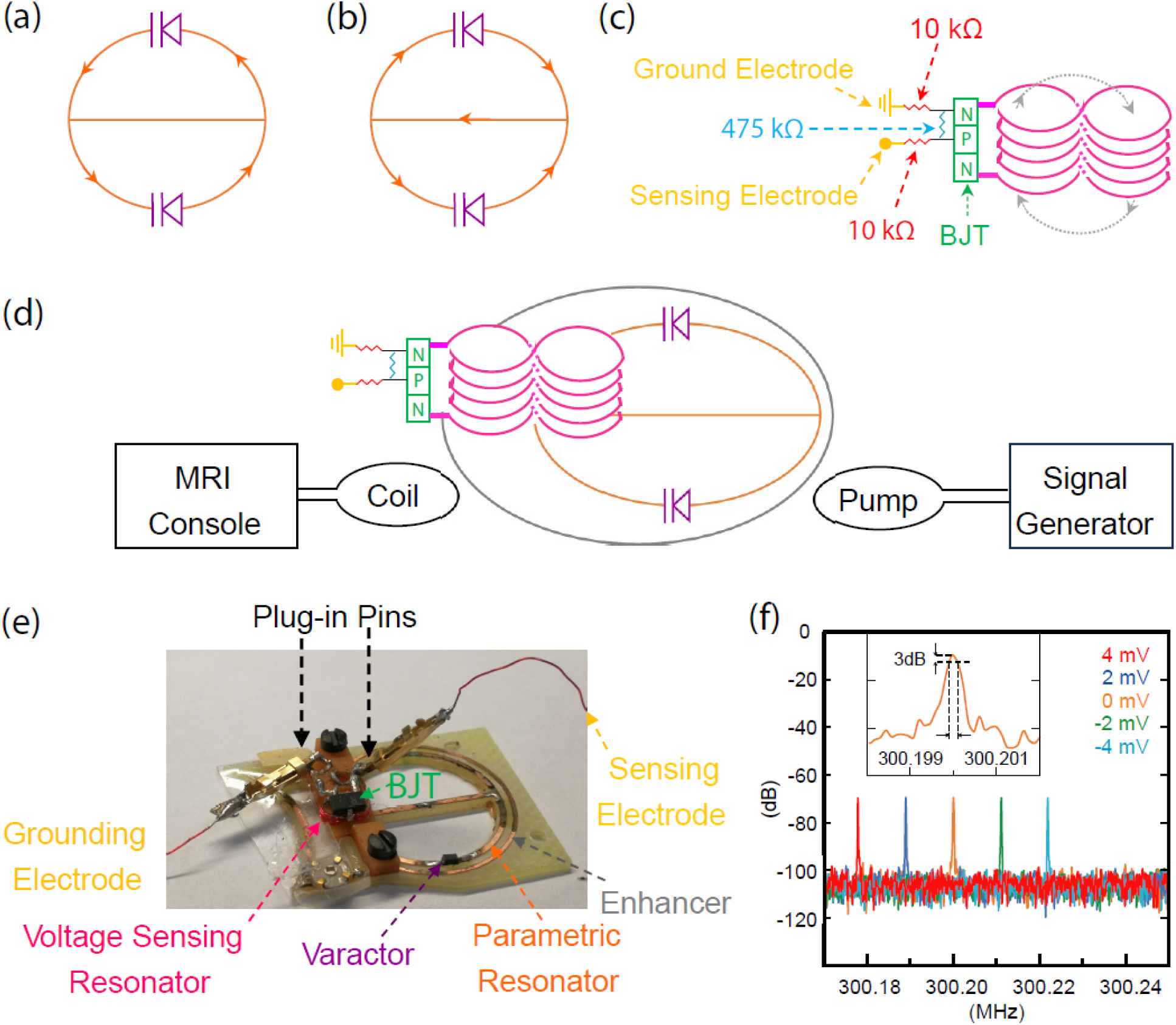
The WISDEM detector consisted of a parametric resonator (a, b) overlaid by a voltage sensing resonator (c). The parametric resonator had a circular shaped conductor pattern (orange) that was split by a pair of varactor diodes (purple) connected in head-to-head configuration, creating a resonance mode with circular-shaped current flow (a). The parametric resonator also had a continuous center conductor to create a second resonance mode with butterfly-shaped current flow (b). The voltage sensing resonator (c) was made by consecutively wrapping an enameled copper wire (pink) around two parallel rods, thus creating five counterclockwise turns in the first rod and another five clockwise turns in the second rod. The wire’s two end edges were soldered to the emitter and collector terminals of a bipolar junction transistor (green). A pair of electrodes (yellow) for sensing and grounding were connected to the transistor’s gate and emitter through 10-kΩ resistors (red). The transistor’s emitter and gate were connected by a 475-kΩ resistor (blue) to provide sufficient internal impedance. (d) The voltage sensing resonator (VSR) was overlapping across the edge of the parametric resonator, with one loop sitting inside the conductor pattern of the Parametric Resonator (PR) and the other loop sitting outside, thus creating effective coupling with the PR’s circular mode resonance. Because both loops were symmetric with respect to the horizontal center conductor of the PR, the voltage sensing resonator (pink) was decoupled from the butterfly mode of the parametric resonator (orange). When the WISDEM was activated by a pumping signal at approximately the sum frequency of the circular and butterfly resonance frequencies, the WISDEM produced sustained oscillation signals for both resonance modes, which could be detected by a standard MRI coil that was cable-connected to the scanner console. (e) When a bias voltage was applied across the pair of electrodes, it linearly shifted the oscillation frequency at a rate of 5.5 kHz/mV (f). This frequency-to-voltage ratio (FVR) was 55-fold larger than the 3dB-linewidth of the oscillation peak (∼100 Hz as shown in the figure insert), enabling sensitive detection of changes in bias voltage as small as 18 µV.

To fabricate a parametric resonator, we first used a CNC milling machine to create a circuit pattern on a copper clad G10 board. This pattern consisted of a circular inductor with an inner diameter of 13.46 mm and an outer diameter of 14.46 mm, leading to an effective inductance of 29.9 nH. Within this circuit pattern, the upper and lower half circles had split gaps that were filled by varactor diodes (BBY53, Infineon, Germany) connected in head-to-head configuration. As a result, the resonator had a resonance mode at 399.5 MHz (Q=79) with circular-shaped current flow (**Fig. 1a**). By connecting the two virtual voltage grounds of the circular mode with a horizontal conductor, a second resonance mode was created at *ω_br_* = 300.2 MHz (Q=77) with butterfly-shaped current flow (**Fig. 1b**). Because the horizontal conductor was connecting the virtual voltage grounds of the circular mode, introduction of the second resonance mode will hardly affect the first resonance mode. To efficiently activate the parametric resonator at the sum frequency of its circular and butterfly modes, pumping field was locally concentrated by an oblong shaped enhancer surrounding the parametric resonator. Fabricated out of a loop conductor with a 15.46-mm width and a 20-mm length, the enhancer was empirically tuned by a trim capacitor that filled its conductor gap. As a result, when the parametric resonator was enclosed inside enhancer, its circular mode resonance frequency was decreased to *ω_cr_*=380.8 MHz (Q=79) while its butterfly mode resonance frequency remained unchanged at 300.2 MHz. Concurrently, the enhancer’s resonance frequency was adjusted to 676 MHz, which was slightly below the sum of butterfly-mode resonance frequency (*ω_br_*=300.2 MHz) and circular-mode resonance frequency (*ω_cr_* =380.8 MHz).

The voltage sensing resonator (VSR) had a figure-8 conductor pattern (**Fig. 1c**). It was fabricated by wrapping a 32-G enameled copper wire around two 1.46-mm diameter rods that were separated by 1.8 mm. Each counterclockwise turn in the first rod was followed by a clockwise turn in the second rod. In this way, five turns with opposite orientations were wrapped around each rod before the wire’s two end terminals were connected to the emitter and collector of a Bipolar Junction Transistor (MT3S111, Toshiba, Japan), creating an effective resonance at 386 MHz (Q=75). Meanwhile, the BJT’s base was connected to its emitter via a 475 kOhm resistor (RC3-0603-4703J, IMS, labelled in cyan in **Fig. 1c**). This 475-kOhm resistor can neutralize excessive charge accumulated on the BJT’s base while maintaining sufficient internal impedance for the transducer. By connecting the BJT’s base with a sensing electrode via a 10-kOhm resistor and the BJT’s emitter with a grounding electrode via another 10-kOhm resistor, the resonance frequency *ω_br_* of the voltage sensing resonator can be effectively modulated by the bias voltage applied across the electrode pair. Meanwhile, the two 10 kOhm resistors (labelled in red) can effectively isolate the entire RF circuit from the sensing electrodes that directly touch biological tissues, thus improving circuit stability. According to the voltage division relation, these two 10-kOhm resistors will only reduce the sensing voltage by a factor of 4% when they are serially connected to the internal impedance of the transducer that is mostly defined by the 475-Ohm resistor between the base and the emitter.

When the voltage sensing resonator was overlapping across the circular edge of the parametric resonator (**Fig. 1d-e**), with one loop sitting inside the parametric resonator and another loop sitting outside the parametric resonator, the VSR could effectively couple with the circular mode of the parametric resonator and decreased the circular mode resonance frequency to 374.8 MHz (Q=67). It is worthwhile to mention that both circles of the voltage sensing resonator were symmetrically distributed across the center conductor line of the parametric resonator, and the VSR’s interaction with the butterfly mode of parametric resonator was effectively cancelled. As a result, the VSR was effectively interacting with only the circular mode of the parametric resonator, enabling effective modulation of the oscillation frequency. When a pumping signal was provided at 675.0 MHz by a loop antenna, the parametric resonator had sustained oscillation current at *ω_b_*=300.2 MHz and *ω_c_*=374.8 MHz. When we varied the DC bias voltage across the pair of sensing electrodes, the oscillation signal shifted at a rate of 5.5 kHz/mV (**Fig. 1f**). This rate of frequency shift was defined as the frequency-to-voltage ratio (FVR). The narrow linewidth (∼100 Hz as shown by the figure insert in **Fig. 1f**) of oscillation peak compared to large voltage-induce frequency shift will enable the wireless detector to identify input voltages as small as 18 µV.

### 2. Retrieving Low-Frequency Voltage Signals Applied on the Sensing Electrodes

To simulate neuronal input signals, we directly injected waveforms produced by a function generator into the sensing electrodes. The function generator produced 20 pulses in every other 1s. Each pulse had a duration of 20 ms, corresponding to one complete sinusoidal cycle. A 10-mm Bruker surface coil was placed behind the WISDEM to relay the oscillation signal into the scanner console (**Fig. 2a**). Once the oscillation signal was recorded, its instantaneous frequency shift was obtained by derivatizing the phase of oscillation signal followed by low pass filtering. Afterwards, the input waveform (**Fig. 2b**) was obtained by dividing this frequency shift with the oscillator’s frequency-to-voltage ratio (FVR=5.5 kHz/mV). **Fig. 2c** showed the retrieved waveform (red) that was averaged over all the five epochs, which agreed well with the input waveform. When we varied the input waveform intensity, we observed a 1:1 linear relation (**Fig. 2d**) between the peak input voltage and the peak voltage value of the reconstructed waveform. This high-level consistency demonstrates reliable voltage encoding capability of the WISDEM.

**Fig. 2.**
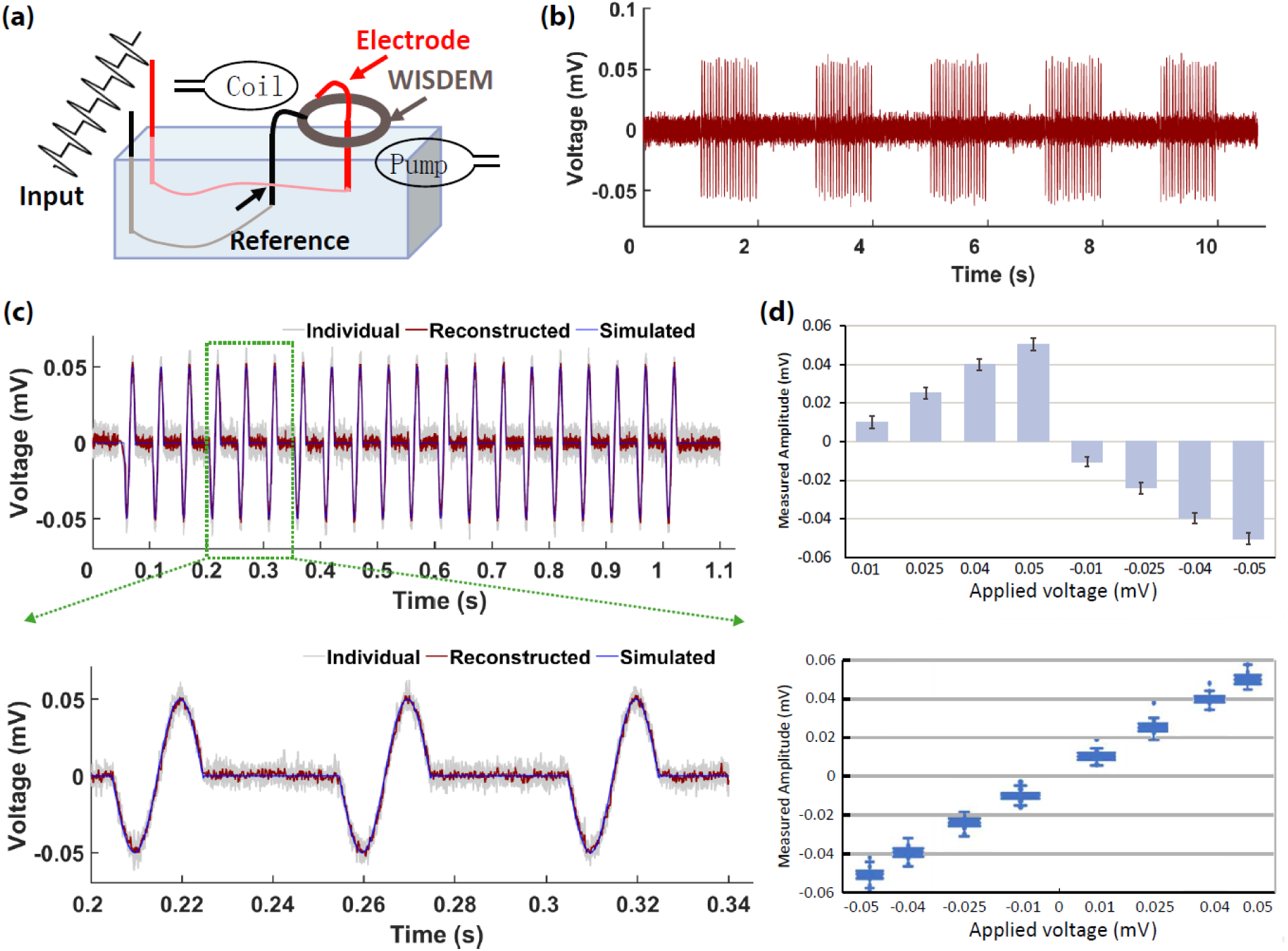
(a) The schematic setup to characterize the frequency response of the WISDEM detector when a train of sinusoidal waves were directly injected into the sensing electrodes. (b) The sinusoidal waveform was reconstructed by derivatizing the phase of oscillation signal and dividing this derivatized value with the frequency-to-voltage ratio, *i.e*. (*d*∅_*t*_/*dt*)/FVR. (c) The zoom-in view of the first epoch containing 20 sinusoidal pulses. When the retrieved waveforms from multiple epochs were stacked together (light gray), their averaged profile (red) was in good agreement with the simulated input waveform (blue), as was clearly demonstrated in the zoom-in insert at the bottom left of the figure. (d) When the peak voltage of the input waveform was systematically varied, it had an obvious linear relation with the peak voltage of the reconstructed waveform. Error bars correspond to Standard Deviation.

### 3. Retrieving High-Frequency MR Signals

To evaluate the detector’s performance for MR signal encoding using Echo Planar Imaging sequence, we placed the detector on an agarose phantom (1% agarose dissolved in distilled water). We then adjusted the pumping power to 0.4 dBm above the oscillation threshold^61^ and continuously acquired the oscillation signal during the entire EPI acquisition period. The horizontal FOV_x_ was enlarged to 3-fold the vertical FOV_y_ so that the spectral window was large enough to include the information-encoding sidebands. By empirically adjusting the pumping frequency, the oscillation signal was adjusted to ∼81.5 kHz above water resonance and aligned to the left quarter location in the frequency domain, thus separating from the image center by ¼ of horizontal FOV_x_ (**Fig. 3a**). In this way, the MR signal had the largest distance separation from its mirror and the aliased mirror. Compared to neuronal voltages that modulated the oscillation signal at <1 kHz speed, MRI signals modulated the oscillation signal at the offset frequency (*e.g*., ∼81.5 kHz), showing up as distinct sidebands in the frequency domain. According to the image reconstruction algorithm described in **Fig. 3a** and **Fig. S6**, we derivatized the phase ∅_*t*_ of the oscillation signal at each time point and assigned the high-pass filtered value of this derivative *d*∅_*t*_/*dt* as the amplitude signal *A_t_* for that particular time point, i.e., *A_t_* =*HPF*(*d*∅_*t*_/*dt*). By multiplying this amplitude signal *A_t_* with the phase term *exp(*-*j*∅_*t*_) of the oscillation signal, a phase-sensitive signal *exp(*-*j*∅_*t*_)∗ *HPF*(*d*∅_*t*_/*dt*) was obtained for that time point. After applying 2D Fourier transformation on phase-sensitive signal series, we could retrieve the image slice shown in **Fig. 3b**. Because the mirrored signal and aliased mirror had opposite phase relations with respect to the MR signal, they appeared as a dispersed pattern after 2D Fourier transformation and were not utilized in subsequent analysis.

**Fig. 3.**
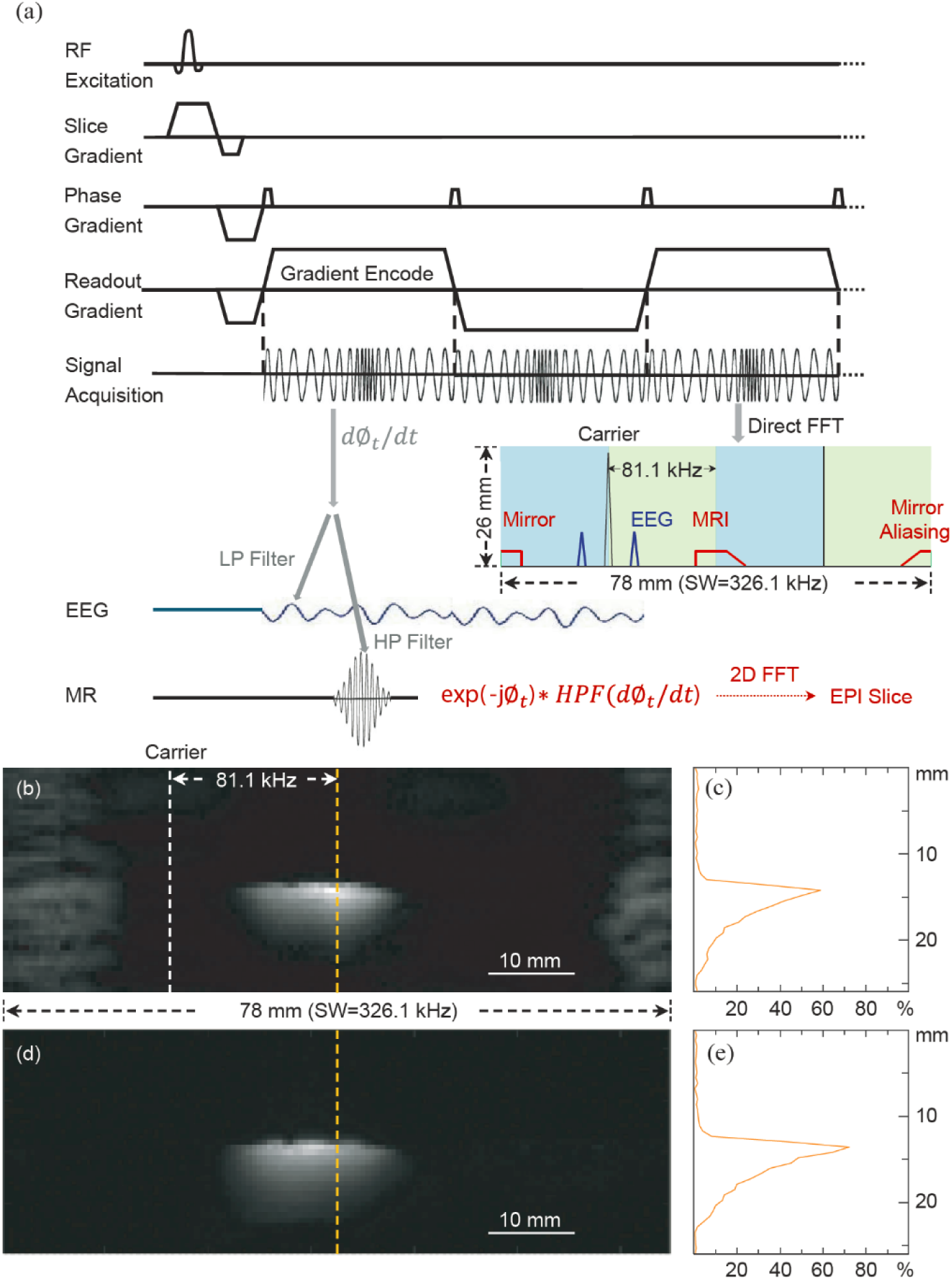
(a) During the gradient encoding periods of Echo Planar Imaging sequence, low-frequency EEG signals and high-frequency MRI signals were simultaneously encoded onto the same oscillation carrier wave that could be directly detected by a standard MRI coil. The oscillation carrier frequency was adjusted to overlap with the left quarter location (represented by the white dash line) of the field-of-view in the frequency domain, so that MR signal was separated from its mirror and the mirror’s aliasing by largest distances. To retrieve time-domain signals, the oscillation carrier wave was first demodulated by derivatizing its phase angle, before being low-pass filtered to obtain EEG and high-pass filtered to obtain MRI. An example of this signal retrieval scheme was shown in Supplemental Fig. S6. (b) A phantom image reconstructed from the oscillation signal recorded during the EPI sequence, using TE = 20.381 ms, TR = 997.648 ms, FOV = 78 × 26 x 14.4 mm^3^, Matrix Size = 129 × 43, Voxel Size = 0.6 × 0.6 mm^2^, Slice Number = 24, Flip Angle = 90°, Bandwidth = 326087 Hz. The dispersed patterns near the left and right edges of the FOV came from the signal mirror and the mirror’s aliasing that had opposite phase relations with respect to the original MR signal in the center. As a result, they were not reconstructed correctly with the correct phase and were discarded for subsequent analysis. (c) Compared to a cable connected coil with identical dimension, the WISDEM maintained ∼60% sensitivity in its sensitivity profile (along the yellow dashed line crossing through the image center). (d) When the pumping power was reduced beneath the circuit’s oscillation threshold, the WISDEM detector could only amplify and relay MRI signals, maintaining ∼75% sensitivity of a cable-connected coil.

To evaluate image sensitivity, we repeated the same procedure to obtain a second image (*S*_2_) and calculated the signal-to-noise ratio (SNR) of individual pixels by dividing the average intensity of individual pixels with the standard deviation of background signal intensity in the difference image.

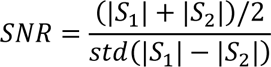

For comparison purposes, we also acquired the same EPI image with a surface coil of the same dimension but with direct wired connection to the scanner console. Compared to this reference image, the image reconstructed from the oscillator maintained ∼60% the sensitivity of a directly connected coil (**Fig. 3c**). Besides full-scope operation as a wireless oscillator, the WISDEM could also operate as a wireless amplifier when the pumping power was reduced to ∼1 dBm below the oscillation threshold. As shown in **Fig. 3d**, when only performing its partial function for signal amplification and wireless transmission^60^, the detector could retain ∼75% the sensitivity of a directly connected coil. Therefore, partial operation as a wireless amplifier would be more suitable for consecutive acquisition of MRI signals during the acquisition intervals of EEG signals, if simultaneous encoding of EEG signals is not required.

### 4. Retrieval of BOLD signals *in vivo* with electrical forepaw stimulation

Next, we demonstrated the detector’s capability of recording BOLD signals *in vivo*. To verify the rat’s brain had hemodynamic responses to sensory stimulation, we first placed the rat inside an MRI scanner and stimulated its somatosensory cortex via its forepaw (**Fig. 4a, b**), using 0.33-ms biphasic electric pulses with 2-mA amplitude at 5-Hz repetition rate that lasted for 4s, followed by a 11s blanking period to restore the activated brain region to its resting state baseline (**Fig.4b**). Concurrently during forepaw stimulation, we activated the WISDEM by a pumping signal at 1 dBm beneath the detector’s oscillation threshold so that we could repetitively acquire Echo Planar Images with enhanced amplitude and good sensitivity. After the amplified images were aligned with the standard SIGMA brain atlas^67^, we calculated the correlation coefficient between the time-dependent signal intensity of each pixel and the ideal stimulation function, thus identifying brain regions that were activated by forepaw stimulation. As shown in the color-coded activation map (**Fig 4c**) overlapped on the atlas^67^, the S1FP region had a BOLD modulation pattern with the same periodicity (15 s) as the stimulation epoch, which was highly consistent with previous results^68–70^ using conventional surface coils. Within each epoch, the MR signal intensity had up to 1.5% modulation during the 4-s stimulation period before returning to the baseline level.

**Fig. 4.**
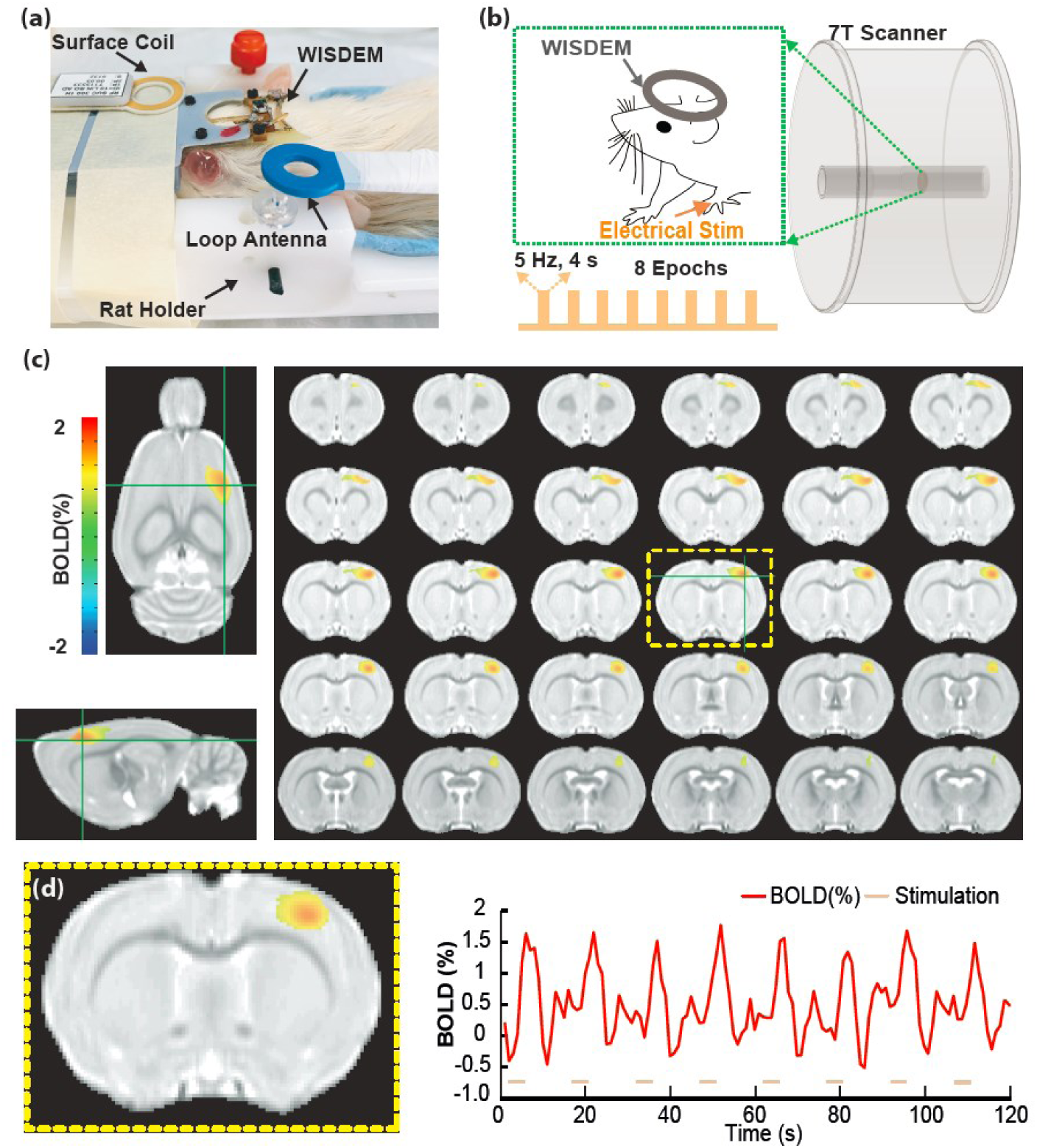
EPI-based functional MRI using the WISDEM from rats upon electrical forepaw stimulation. (a) The picture of a WISDEM detector placed on top of a rat head that was fixed inside a cradle. (b) The schematic diagram showed the rat secured inside the MRI scanner, with its left forepaw stimulated by 8 epochs of electric pulses. Each epoch contained 333-µs biphasic pulses at 5 Hz repetition rate and 4-s duration followed by 11-s resting period. (c) When the WISDEM was operating as an MR signal amplifier, the evoked BOLD fMRI maps showed a clear activation region in the forelimb area of the primary somatosensory cortex (S1FP) based on the SIGMA rat brain atlas^67^, following tactile stimulation of the left forepaw (*n* = 5 animals, *P*_(corrected)_ < 0.001). (d) The modulated signal pattern in the S1FP region was highly synchronized with the stimulation pattern (shown as pink dash) applied on the rat’s left forepaw.

### 5. Simultaneous Retrieval of BOLD and LFP signals *in vivo* with optogenetic stimulation

To demonstrate the full-scope capability of the WISDEM, we simultaneously recorded LFP during MR signal acquisition. Rats were stimulated by light pulses at 470 nm wavelength, with an optical fiber and an electrode inserted into the S1FP region that had been transfected with AAV5-CaMKII.hChR2 (**Fig. 5a, b**). To confirm the sensing electrode was recording LFP signals, we acquired the detector’s oscillation signals in the absence of encoding gradients or RF pulses when light stimulation was repetitively applied at 2 Hz for 3 pulses followed by a 1s rest interval. Then, we derivatized the phase angle of the oscillation signal *d*∅_*t*_/*dt* to obtain time-dependent electrophysiological signals. As shown in the upper row of **Fig. 5c**, EEG spikes were clearly observable for low-power light at 0.39 mW. In a control experiment, we reduced light power to 0.005 mW and observed disappeared LFP patterns (bottom row of **Fig. 5c**), thus attributing the negative peaks in the upper figure to light induced activity. EEG spikes were robustly recorded with laser power dependency (0.39, 0.97, 1.58, 2.20, 2.83 mW in **Fig. S1**) and laser width dependency (1, 5, 10, 15, 20 ms in **Fig. S2**). When we increased the laser power, we observed decreased latency time for the negative peak (**Fig. 5d**). When optical stimulation power was increased, the decreased latency time was also observed for the positive peak that always followed the negative peak in the EEG curve. No LFP spikes were observed when the same amount of maximum laser power was applied on a control rat without AAV-ChR2-mCherry expression (**Fig. S4**), thus once again attributing the negative peaks in **Fig. S1-S2** to optogenetically evoked activation. Meanwhile, the increased stimulation power also led to larger peak intensity (**Fig. 5e**, n = 4 animals). It is noteworthy that the orange curve for “rat 2” had larger deviation from the curves corresponding to the other three rats. This is because the electrode was inserted shallower than the virus injection point to verify the spatial specificity of optogenetic stimulation and the longer duration for neuronal currents to travel over a larger distance separation. To confirm the rat’s alertness, we also used the same method to retrieve EEG signals when the rat was stimulated by electrical currents on its forepaw (**Fig. S5**).

**Fig. 5.**
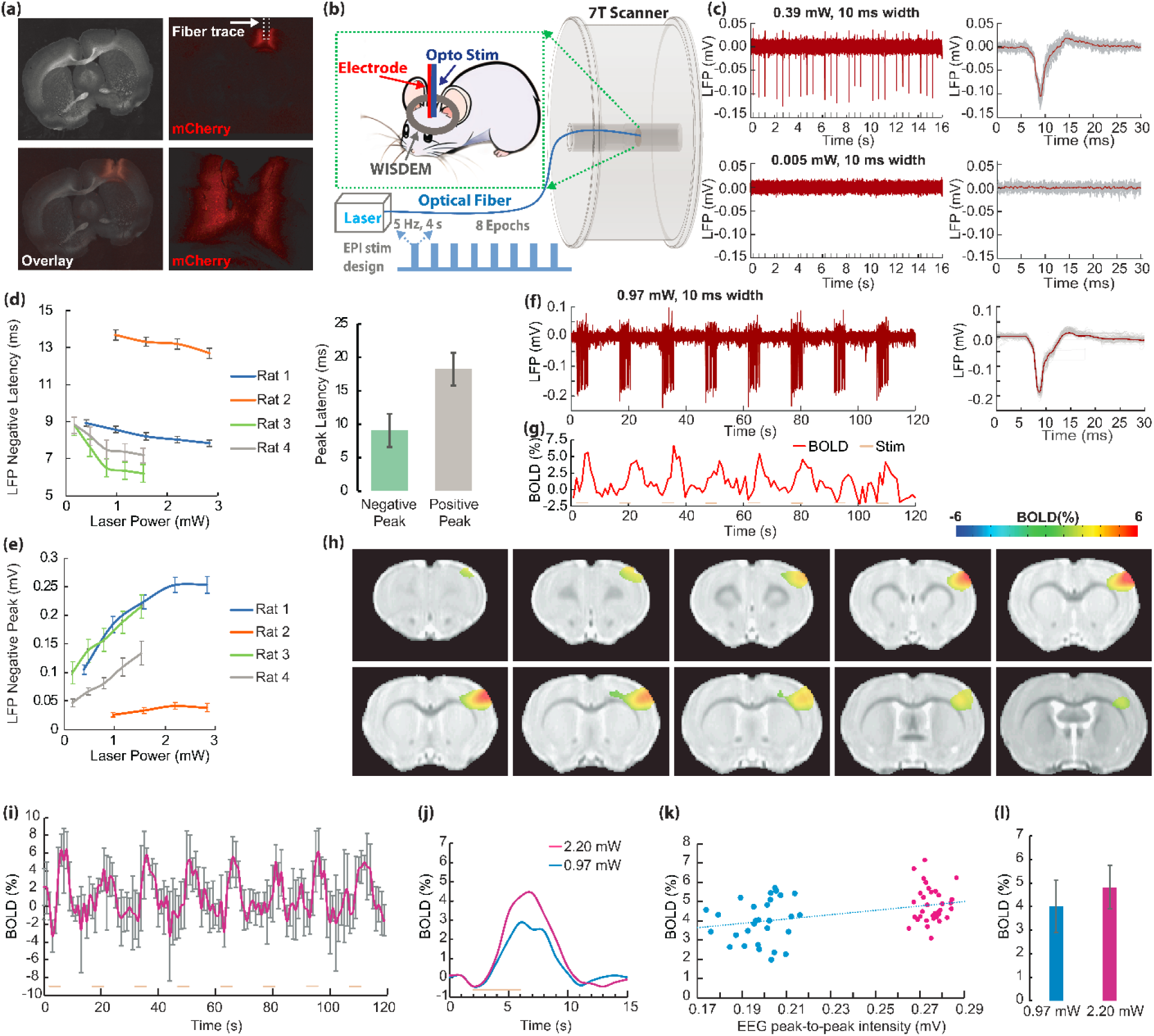
WISDEM for optogenetic-fMRI in rats. (a) The histology brain slice and fluorescence images showed the fiber injection spot in the S1FP region with AAV-ChR2 expression. (b) The schematic diagram showing the rat lying inside the MRI scanner, whose S1FP region was stimulated by an optical fiber and recorded by a sensing electrode. (c) In the absence of encoding gradients, the EEG pattern could be directly retrieved when the stimulation light power was as low as 0.39 mW. The grey lines in the right column showed all the EEG traces after individual stimulation pulses, while the red line represents the average value of gray traces. (d) When the laser power gradually increased, AAV-labelled neurons had stronger responses, leading to decreased latency time and (e) increased peak amplitude. (f) In the presence of encoding gradients, EPI sequence was implemented concurrently during optogenetic stimulation. The EEG signal was retrieved by low pass filtering the derivatized phase angle of the detector’s oscillation signal, *LPF*(*d*∅_*t*_/*dt*). (g) To retrieve multi-slice MR images, we implemented the signal reconstruction algorithm described in Fig. 3a and Fig. S6. The S1FP region showed a modulated signal pattern that was highly synchronized with the stimulation epoch and the EEG pattern. (h) The color-coded activation map depicted regions with synchronized intensity modulation that were overlapped on the SIGMA rat brain atlas^67^ of Block design (Single-sided GLM-based t-statistics in AFNI was used. *P* (corrected) < 0.001). (i) Averaged BOLD fMRI time courses from the activation region in S1FP upon optogenetic stimulation (n = 4 animals, mean ± SD). (j) Averaged BOLD response for each epoch with different light intensities. (k) The EEG peak intensity and fMRI signal response show positive correlation for individual epochs. The blue and magenta dots correspond to laser power at 0.97 mW and 2.2 mW, respectively. (l) The BOLD peak responses under two different laser power levels. Error bars correspond to Standard Deviation.

Simultaneously, we repetitively executed Echo Planar Imaging sequence (in the presence of encoding gradients and RF pulses) and concurrently recorded the oscillation signal during optogenetic stimulation. Using the reconstruction algorithm described in **Fig. 3a** and **Fig. S6**, we could simultaneously retrieve EEG and fMRI signals. When the optical stimulation power was 0.97-mW, the LFP pattern (**Fig. 5f**) had most peaks retaining similar amplitudes as those acquired in the absence of gradients or RF pulses (**Fig. S1b**). As shown in Supplemental **Fig. S3**, when the optical power level was increased from 0.97 mW to 2.20 mW, the LFP peaks retrieved in the presence and in the absence of RF pulses & encoding gradients maintained to have comparable intensity, thus demonstrating the robustness of the wireless oscillator for LFP signal encoding during MR signal acquisition. In the concomitantly acquired EPI image, the S1FP region had a periodically modulated pattern with an average of ∼3% intensity increase during the 4-s stimulation period of 0.97 mW pulses (**Fig. 5g**), which was highly consistent with previous studies^71–73^ using conventional surface coils. This fMRI signal pattern was also synchronized with the LFP pattern (**Fig. 5f**). Based on the mask defined in the color-coded activation map (**Fig. 5h**), we evaluated the signal intensity over multiple EPI experiments and consistently obtained a similar time-dependent fMRI modulation pattern (**Fig. 5i**). By plotting the fMRI signal change during each stimulation epoch with respect to the corresponding peak LFP intensity, we obtained an approximate linear relation between fMRI intensity change and LFP intensity (**Fig. 5k**), thus demonstrating the presence of neurovascular coupling effect that led to MRI signal modulation.

## Discussion

In this work, we have created a WISDEM platform that combines, for the first time, LFP and BOLD recording through the MR console system without the need for extra recording equipment. The 2-in-1 WIDEM detector can wirelessly communicate with any type of signal interface that is already available on commercial MRI scanners, thus providing an easy-to-access-tool to interrogate neuro-vascular coupling mechanisms in healthy and diseased brains.

Our WISDEM transducer has a very compact design. Without the need for dedicated signal amplifiers, the WISDEM is a high-quality oscillator that is very sensitive to small input signals, enabling simultaneous encoding of both low-frequency EEG signals and high-frequency MRI signals onto the same oscillation carrier wave. The encoded signals appear as distinct sidebands that are easily separable in the frequency domain. Because EEG and MRI signals are retrieved from the same wireless carrier wave that can be detected by a standard MRI coil, no dedicated hardware is required to synchronize these two detection modalities. The WISDEM transducer consists of two nonlinear circuits that can be individually optimized, *i.e.*, the Voltage Sensing Resonator (VSR) for low-frequency signal encoding and the Parametric Resonator (PR) for high-frequency signal encoding and wireless carrier broadcasting. The PR has a circular mode and a butterfly mode to sustain oscillating current flows. The PR can utilize wireless pumping power provided at the sum frequency of its two resonance modes and the multi-band frequency mixing process to provide power for circuit oscillation at the circular and butterfly modes. On the other hand, the voltage sensing resonator is made by connecting the emitter and collector terminals of a Bipolar Junction Transistor with an 8-shaped conductor wire. Unlike the previous design^66^ that used two varactor diodes in head-to-head configuration, the voltage sensing resonator used here has an inductor connected to a single-element transistor. Since only two soldering junctions instead of four are required to complete the resonance circuit, the VSR has a higher quality factor (*Q*=75) with smaller parasitic resistance. When the neuronal voltage applied on the transistor’s base varies over time, the emitter-base junction capacitance *C_eb_* and the collector-base junction capacitance *C_cb_* are varied at the same pace, leading to effective modulation of the VSR’s resonance frequency. Also, because the transistor’s terminals are connected to sensing electrodes via buffering resistors, radio-frequency noises from biological tissues are mostly blocked by the buffering resistors, thus minimizing circuit loss. When the VSR couples to the circular resonance mode of the PR, resonance frequency shift of the VSR can be efficiently converted into oscillation frequency shift of the PR, and the time-dependent oscillation signal can be wirelessly detected by a standard MRI coil with cable connection to the scanner console. Because both the PR and VSR have low circuit loss, the oscillation signal of the entire WISDEM transducer has a narrow linewidth (100 Hz), enabling efficient detection of low-frequency voltage signals as small as 18 µV. For high-frequency MRI signals, the PR can combine down conversion and frequency encoding into a single stage, thus modulating the oscillation signal at the offset frequency between the input signal and the oscillation signal. After derivatizing the phase angles of the oscillation signal, the MR signal corresponding to individual time points can be retrieved by high-pass filtering. Image reconstruction by 2D Fourier Transform can be correctly performed when each magnitude term *HPF*(*d*∅_*t*_/*dt*) is multiplied by the phase factor *exp(*-*j* ∅_*t*_ ) of the oscillation signal.

Another advantage of the WISDEM is the transducer’s signal encoding capability is minimally perturbed by imaging sequences. This is because MRI signals are received by the butterfly mode of the parametric resonator that is tuned approximately to the proton Larmor frequency (at 7T). Because the butterfly mode of the Parametric Resonator (PR) has a magnetic field pattern that is perpendicular to the B_1_ field produced by the volume coil, the excitation B_1_ field from the volume coil has minimal interfering effect on the PR. This design averts the need for specially designed electrodes or recording instruments that were traditionally required to minimize noise contamination inside MRI scanners^74,75^. Unlike previous work on simultaneous fMRI and EEG that required graphene electrodes^20^ to reduce electromagnetic artifacts propagating along connection cables, the WISDEM can interface with ordinary metallic electrodes that are widely used in neuroscience labs. Because the WISDEM’s oscillation signal is continuously recorded over the entire duration of MR acquisition windows even in the presence of rapidly switching gradients, fMRI and EEG signals can be reliably retrieved using the simple method described in **Fig. 3** and **Fig. S6**. These desirable features make it convenient to perform synchronous measurement of neuronal and hemodynamic activities in both healthy and diseased animals for mechanistic studies of neurovascular coupling/decoupling.

In addition to observing focal regions that are closer to the brain surface, there are also demands to monitor neuronal activation in deep-lying regions across the entire brain. In our prototype device, the parametric resonator has a diameter of 13.5 mm, creating a butterfly mode with an effective detection depth of ∼10 mm that is already comparable to the radius of a rat brain. To further enlarge the detector’s effective depth, we can potentially utilize the circular mode of the parametric resonator to receive MR signals from deeper regions and align the detector’s normal axis approximately perpendicular to the B_1_ field of a linear-mode volume coil that will be utilized for nuclear spin excitation. Such an arrangement can fully utilize the detection depth of a circular-mode detector and at the same time minimize interfering interactions from the MR excitation pulses, leaving the residual interference easily removable by the baseline correction algorithm described in **Fig. S6**.

To record neuronal activity from different brain layers, it is possible to detect individual layers by multiple electrodes and connect all electrodes to the same Voltage Sensing Resonator via a multiplexer^76–78^. As a result, the same detector can consecutively encode neuronal voltages from multiple electrodes while continuously encoding MRI signals from the same field-of-view. To simultaneously detect multiple brain regions over an extended FOV, it is also possible to construct an array of WISDEM detectors, each of which is wirelessly activated by a unique pumping frequency^79^, enabling independent manipulation of individual detectors.

We have demonstrated an immediate application of the WISDEM detector to characterize neuronal responses during optogenetic stimulation that has improved cellular specificity and reduced electrophysiological noises when compared to electric stimulation. Without the need for connection cables, the WISDEM detector offers extra degrees of freedom for electrode placement, saving precious space for optical fibers that are normally required to deliver light into the brain. A next-stage preclinical application is to incorporate photometry into the optical pathway, thus providing concurrent capabilities for fluorescent calcium recordings and electrophysiology during optogenetic stimulation^80^. For potential clinical applications, this 2-in-1 wireless detector will be particularly advantageous in epileptic patients who already need implantable electrodes, where the simultaneous imaging capability will help better locate disease foci.

In conclusion, we have fabricated a wirelessly powered oscillator that can encode both low-frequency and high-frequency signals for simultaneous EEG and fMRI. Without the need for cable connection to a separate EEG apparatus, this hybrid 2-in-1 transducer can be easily implemented by the neuroscience community, enabling real-time monitoring of neurovascular coupling events underlying fMRI signal dynamics, thus significantly boosting fMRI’s capability as a research and diagnostic tool.

## Acknowledgments

We thank Dr. Yuan Gao for providing technical support for immunohistology. This research was supported by National Institutes of Health (RF1NS113278-01, RF1NS128611-01 to X.Y. and C.Q.) and by the Division of Electrical, Communications and Cyber Systems of the National Science Foundation under award number 2144138 (to C.Q.). This project has received funding from the European Union Framework Program for Research and Innovation Horizon 2020 (2014–2020) under the Marie Skłodowska-Curie Grant Agreement No. 896245. Any opinions, findings, conclusions, or recommendations expressed in this material are those of the author(s) and do not necessarily reflect the views of the funding agencies.

## Author Contributions

X.Y. and C.Q. conceptualized the design concept and acquired funding, Y.C., W.Q. and C.Q. performed experiments, Y.C. and C.Q. wrote the manuscript, D.R. and X.Y. provided technical support, revised the manuscript.

## Inclusion & ethics statement

All collaborators of this study have fulfilled the criteria for authorship required by Nature Portfolio journals have been included as authors, as their participation was essential for the design and implementation of the study. Roles and responsibilities were agreed among collaborators ahead of the research. The authors declare no competing interests.

## Data and Resource Availability

All data needed to evaluate the conclusions in the paper are available on https://doi.org/10.6084/m9.figshare.26082115. The WISDEM device can be provided by the corresponding author pending scientific review and a completed material transfer agreement.

## Code Availability

Codes for converting oscillation signal into MRI and EEG signals were written in Matlab2023a platform. These codes can be downloaded from https://codeocean.com/capsule/6663434/tree. The Analysis of Functional NeuroImages software (AFNI, NIH, USA) was used to process the reconstructed fMRI images. The relevant source codes can be downloaded through https://afni.nimh.nih.gov/afni or provided upon direct email request to the corresponding author.

## Supplementary Information

### Appendix

The parametric resonator requires an externally provided pumping signal to oscillate. Normally, the pumping frequency *ω_p_* can be experimentally adjusted by tuning the knob of an external frequency synthesizer. Once *ω_p_* is set, it will also determine the sum of butterfly and circular mode oscillation frequencies, i.e., *ω_p_ = ω_b_+ ω_c_*. The exact values of *ω_b_* and *ω_c_* can be derived from the following equation that describes the equal relation of reactance-to-resistance ratio in both resonance modes:

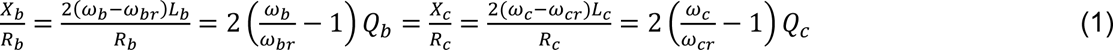

In Eq. (1), *L_b_* and *L_c_* are effective inductance of the butterfly and circular modes, while *R_b_* and *R_c_* are effective resistance of the butterfly and circular modes. By plugging *ω_b_ = ω_p_* − *ω_c_* into Eq. (1), the butterfly mode oscillation frequency can be calculated as:

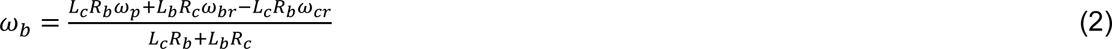

If a voltage sensing resonator can somehow modulate the circular mode resonance frequency *ω_cr_* without affecting the butterfly mode resonance frequency *ω_br_*, the value of *ω_b_* can also be effectively modulated.

### Materials and Methods

#### Animals

All procedures in this study were conducted in accordance with guidelines set by the Institutional Animal Care and Use Committee of Michigan State University. In total 10 Sprague Dawley rats (5 for forepaw stimulation experiments, 4 for optogenetic-fMRI experiments and 1 rat for control experiment without AAV-ChR2-mCherry) from Charles River were used in this study. All animals were three-in-one-housed in 12-12 hour on/off light-dark cycle conditions to assure undisturbed circadian rhythm and ad libitum access to chow and water.

#### Optogenetic virus injection

AAV5.CaMKII.hChR2 (H134R)-mCherry was purchased from Addgene and packaged into frozen-stored vials, each of which contained 100 µL of sample at a concentration higher than 1×10¹³ vg/mL. To inject AAV5 into the right somatosensory forepaw region of brain in a 4-week-old rat, the rat was first anesthetized with 1.5–2% isoflurane via a nose cone and secured on a stereotaxic frame. An incision was made on the scalp to expose the skull before craniotomies were made with a pneumatic drill to introduce minimal damage to cortical tissue. Afterwards, 0.4–0.6 µL of viral droplet was injected from a 10-µL syringe via a 35-gauge needle to the following coordinates: 0 mm posterior to the Bregma, 3.2–3.5 mm lateral to the midline, 0.5–1.2 mm below the cortical surface using an infusion pump (Pump 11 Elite, Harvard Apparatus, USA). Once AAV5 injection was finished, the needle was left in place for approximately 5 min before being slowly pulled out. The craniotomies were sealed with bone wax, and the skin around the wound was sutured. After the surgery, rats were subcutaneously injected with antibiotics and painkillers (Ketoprofen fluids) for three consecutive days to prevent infection and to relieve pain. Imaging experiments were performed 4 weeks after virus injection to allow enough ChR2 protein expression in the S1FP region.

#### Animal preparation, electrode and fiber implantation

To prepare for electrode implantation, an enameled copper wire of 80-µm diameter was glued with an optical fiber of 200-µm core diameter (FT200EMT, Thorlabs). The front edge of the copper wire was trimmed by a scissor to expose the conductor tip for direct contact with the brain tissue. Once the animal was anesthetized by 2% isoflurane with its head fixed on a stereotaxic frame, a burr hole of 1.5-mm diameter was drilled on the rat’s skull so that the dura can be carefully removed. Afterwards, the optical fiber along with the attached electrode was inserted into the S1FP region, at coordinates of 0 mm posterior to Bregma, 3.2-3.5 mm lateral to the midline and 1.2 mm below the cortical surface. Subsequently, an adhesive gel (Loctite 454, Henkel, Germany) was applied on the insertion hole to secure the fiber/electrode assembly against the skull. A grounding electrode was separately screwed against the skull above the neck. After the scalp was closed by glue, the rat was injected by a bolus of dexmedetomidine (0.05 mg/kg, sc; Dexdomitor®, Orion Pharma) before isoflurane was discontinued. The rat was then transferred into the MR scanner with its head secured by a bite bar and two ear bars, so that the WISDEM detector could be mounted above the rat’s head to cover the right S1FP region (Fig. 4a). Finally, both the sensing electrode and the grounding electrode were connected to the plug-in pins that were previously soldered on the voltage sensing resonator.

During *in vivo* imaging, the rat was subcutaneously administered with constant infusion of dexmedetomidine at 0.1 mg/kg/hr. Its breathing and heart rates were monitored by an air pillow placed beneath its chest that was interfaced with an MR-compatible monitoring system (Model 1025, SA Instruments, Inc., Stony Brook, NY). The rat’s body temperature was continuously monitored by a rectal probe (SA instrument) and maintained at 37°C by a water jacket.

#### Functional MRI acquisition

All MRI data were acquired inside a 7T small-animal scanner (Bruker BioSpin, Billerica, MA) with a 16-cm horizontal bore. The WISDEM detector was placed above the animal’s skull and activated by a loop antenna to produce sustained oscillation signal. Functional MR images were acquired with a multi-slice gradient-echo EPI (GE-EPI) sequence with the following parameters: time of echo (TE) = 20.381 ms, time of repetition (TR) = 1 s, field of view (FOV) = 78 × 26 x 14.4 mm^3^, matrix size = 129 × 43, voxel size = 0.6 × 0.6 mm^2^, slice number = 24, flip angle = 90°, bandwidth = 326087 Hz. Concurrently during image acquisition, we also performed electrical stimulation on the rat’s left forepaw or optogenetic stimulation in the S1FP region. For both the forepaw and optogenetic stimulation, the stimulation paradigm started from a 11-s pre-stimulation delay, followed by 8 epochs of stimulation cycles, each of which starts from a 2-s resting period followed by 4-s stimulation period and concludes by a 9-s interval. As a result, the total repetition number for the entire EPI experiment was 135. The 4-s period for electrical forepaw stimulation contained 20 biphasic pulses, each of which had 333-μs duration and 5-Hz repetition rate. The 4-s period for optogenetic stimulation contained 20 square pulses, each of which had a 10-ms duration and 5-Hz repetition rate.

#### MRI data analysis

After functional images were retrieved using the algorithm described in Figs. 3a and S6, they were imported into the AFNI software (Analysis of Functional NeuroImages, NIH, USA) for subsequent analysis. First, functional images were aligned to anatomical images that were separately acquired with RARE (Rapid Acquisition with Relaxation Enhancement) sequence. These anatomical images had the same orientation and FOV as functional images but with a higher spatial resolution at 100 µm. The anatomical images were then registered to SIGMA rat brain template^67^ to derive the transformation relation, based on which the functional EPI images were registered to the same standard template. The baseline signal of EPI images was normalized to 100 for statistical analysis of multiple trials over the time course. The time courses of the BOLD signal were extracted from the S1FP region because this region had significant activation values in the brain atlas. The hemodynamic response function (*HRF*) used the *BLOCK* function in the linear program *3dDeconvolve*, where the statement *BLOCK (L, 1)* was a convolution of a square wave of duration *L* and peak amplitude of 1. To compute the evoked BOLD changes in **Figs. 4d, 5h**, *3dmaskave* was evaluated in the ROI defined as the primary somatosensory area in the SIGMA atlas^67^.

#### Immunohistochemistry

To verify the phenotype of the transfected cells, opsin localization, and optical fiber placement, the rat was sacrificed and perfused in its left ventricle. The rat brain was extracted, fixed overnight in 4% paraformaldehyde and then equilibrated overnight in 15% sucrose dissolved in 0.1 M phosphate buffer at 4°C, before being soaked inside 30% sucrose dissolved in 0.1 M phosphate buffer. Subsequently, the brain was sectioned to 30-µm slices on a sliding microtome (Leica CM 1850, Germany). Free-floating brain slices were washed in PBS, mounted on microscope slides, and incubated with DAPI (Sigma Aldrich, USA) at room temperature, before being imaged by a fluorescence microscope (Nikon A1 Laser Scanning Confocal Microscope, Japan) for assessment of ChR2 expression in the S1FP region (**Fig.5a**). To enhance brightness and contrast for visualization purposes, digital images were minimally processed using ImageJ.

---

**Supplemental Fig. 1.**
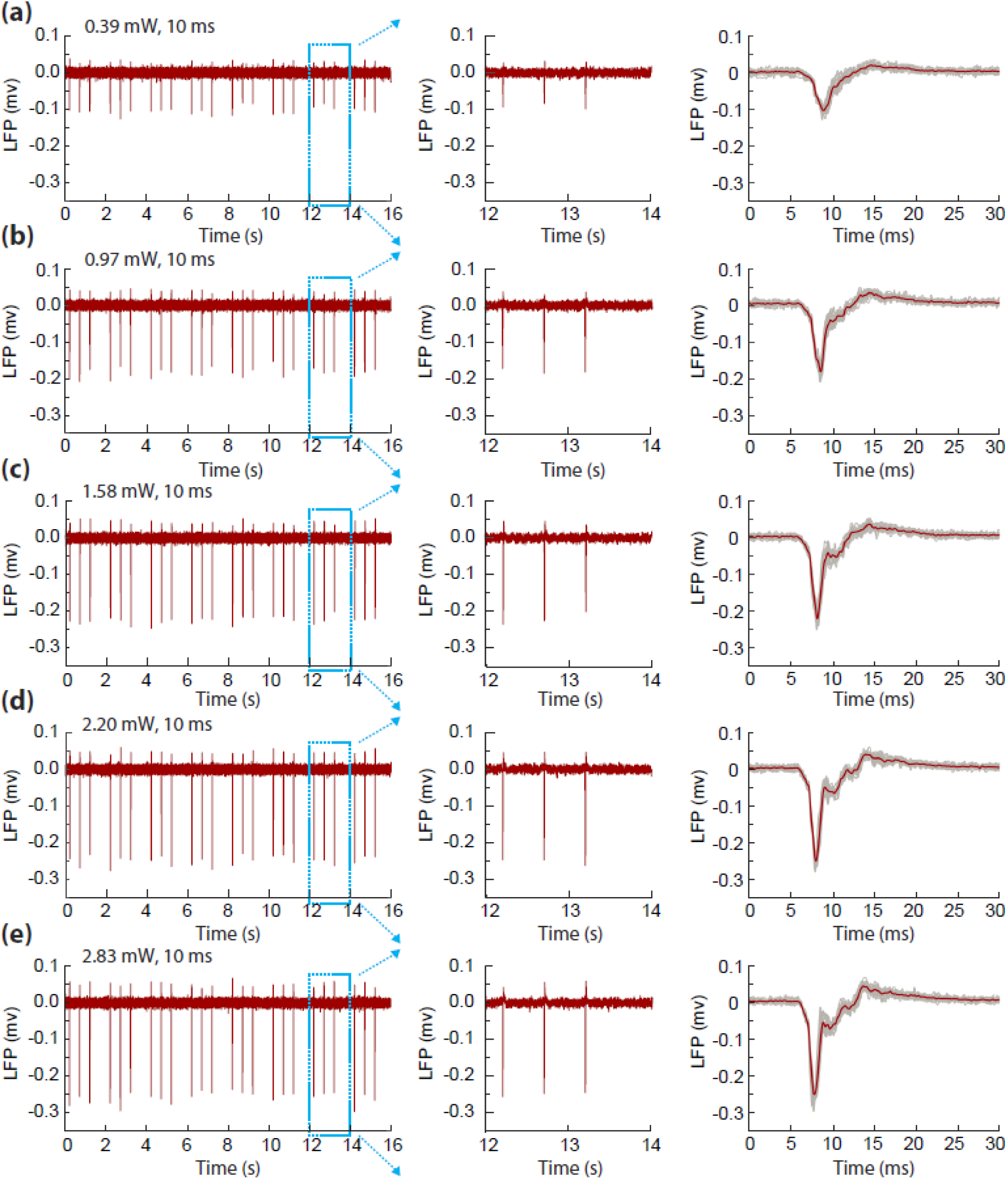
The reconstructed LFP traces obtained during optogenetic stimulation of a representative rat under different light power (a-e). Stimulation pulses were applied at 0.2, 0.4, 0.6, 2.2, 2.4, 2.6, 4.2, 4.4, 4.6, 6.2, 6.4, 6.6, 8.2, 8.4, 8.6, 10.2, 10.4, 10.6, 12.2, 12.4, 12.6, 14.2, 14.4 and 14.6 s following the initial starting point of pulse sequence. The grey lines in the right column show all the LFP traces after individual stimulation pulses, while the red line represents the average value of gray traces.

**Supplemental Fig. 2.**
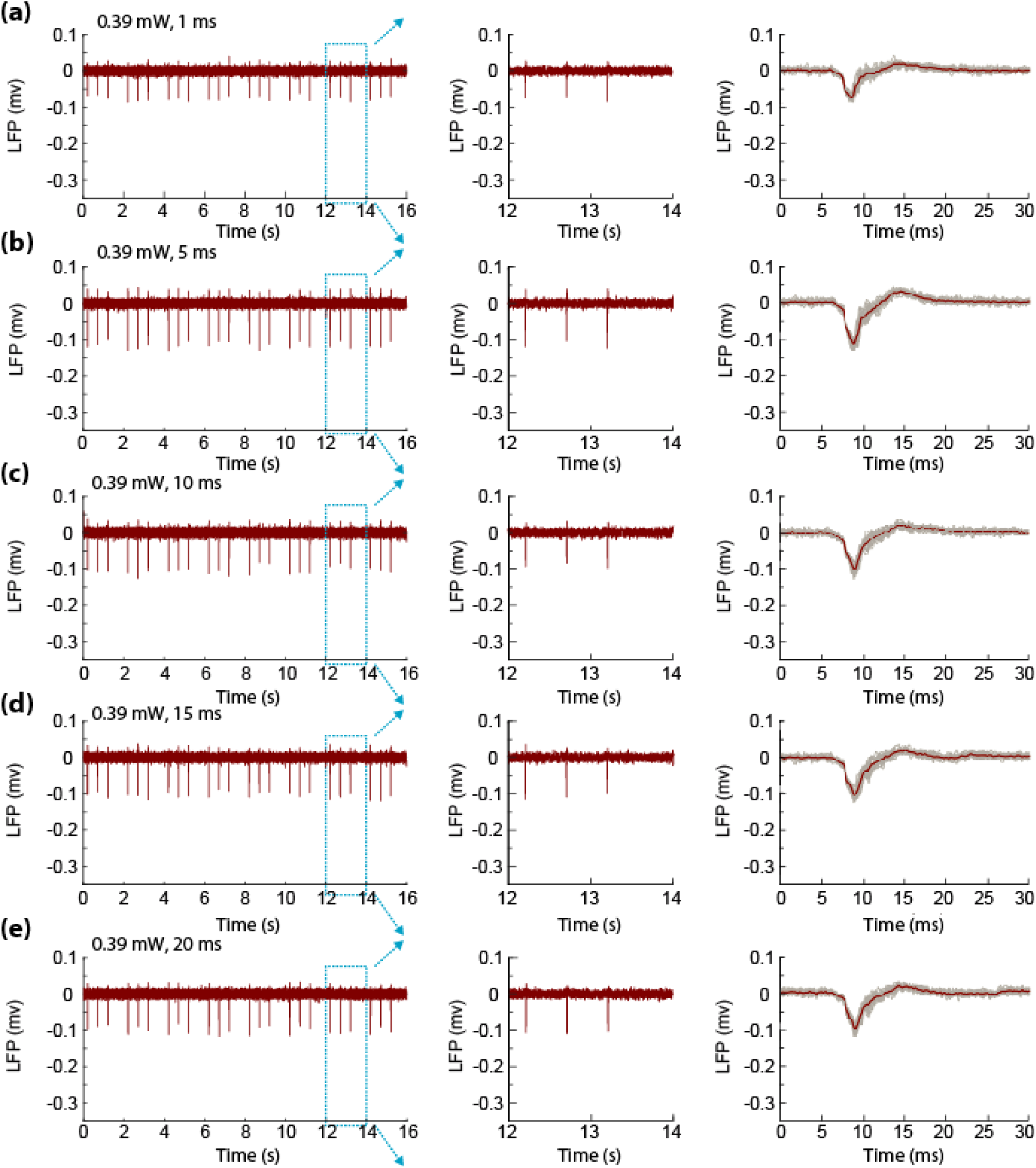
The reconstructed LFP traces obtained during optogenetic stimulation of a representative rat using different light widths (a-e). Stimulation pulses were applied at 0.2, 0.4, 0.6, 2.2, 2.4, 2.6, 4.2, 4.4, 4.6, 6.2, 6.4, 6.6, 8.2, 8.4, 8.6, 10.2, 10.4, 10.6, 12.2, 12.4, 12.6, 14.2, 14.4, 14.6 s. The grey lines in the right panel show all the LFP traces after individual stimulation pulses, while the red line represents the average value of gray traces.

**Supplemental Fig. 3.**
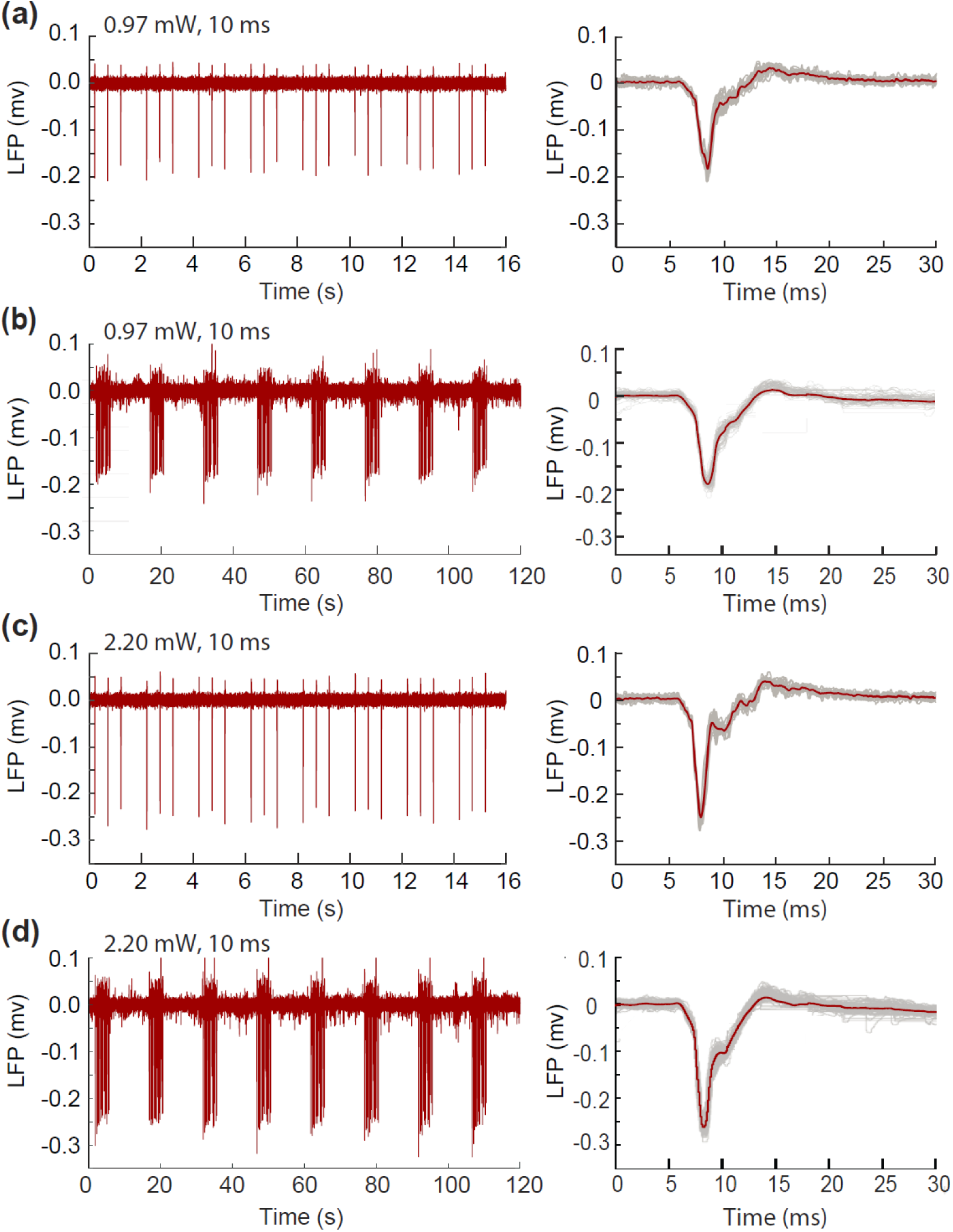
The reconstructed LFP traces obtained during optogenetic stimulation of a representative rat under different light power, showing the comparable peak intensity obtained in the absence (a,c) and presence (b,d) of RF and gradient pulses. When no RF or gradient pulses were present, the S1FP region was stimulated at 2-Hz every other second within a 15-s period to quickly verify the brain was in an arousal state to produce LFP signals. Subsequently, RF and gradient pulses were turned on for Echo Planar Imaging, while the S1FP region was stimulated by 8 epochs of stimulation cycles, each of which started from a 2-s resting period followed by a 4-s stimulation period containing 20 pulses applied at 5 Hz. Each stimulation cycle finally concluded by a 9-s interval. The grey lines in the right panel show all individual LFP from the eight epochs, while the red line represents the average value of gray traces.

**Supplemental Fig. 4.**
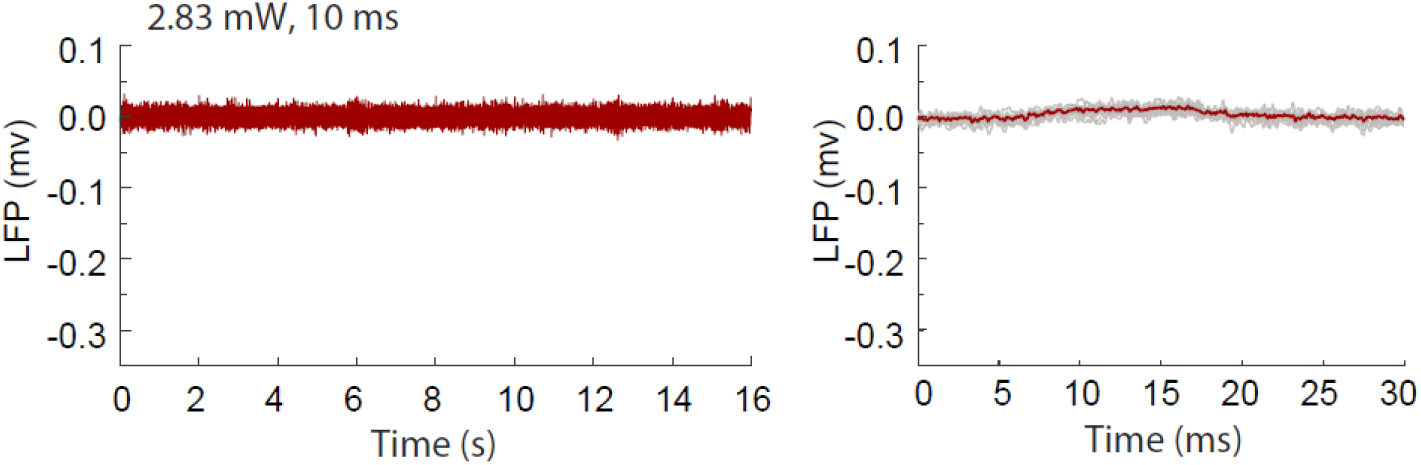
When the same amount of maximum laser power was applied on a control rat without AAV-ChR2 expression, no LFP spikes were observed from the S1FP region, thus demonstrating the neuronal origin of LFP spikes in Figs.S1-3, rather than photoelectric effect.

**Supplemental Fig. 5.**
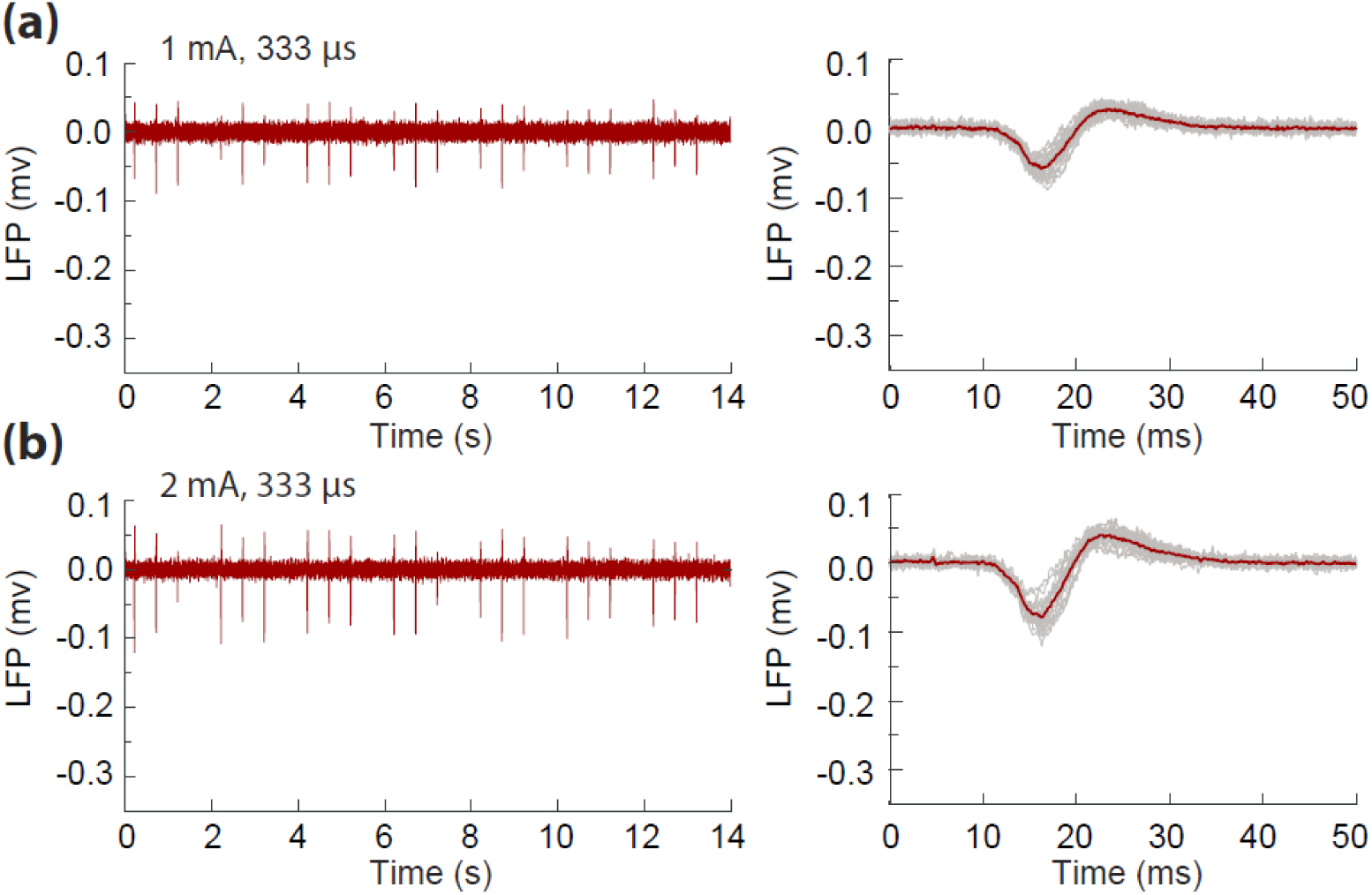
The reconstructed LFP traces obtained during forepaw stimulation of a representative rat under 1 mA (a) and 2 mA (b). Individual stimulation pulses had widths of 333-µs. They were applied at 0.2, 0.4, 0.6, 2.2, 2.4, 2.6, 4.2, 4.4, 4.6, 6.2, 6.4, 6.6, 8.2, 8.4, 8.6, 10.2, 10.4, 10.6, 12.2, 12.4, 12.6s. The grey lines in the right column show all the LFP traces after individual stimulation pulses, while the red line represents the average value of gray traces.

**Supplemental Fig. 6.**
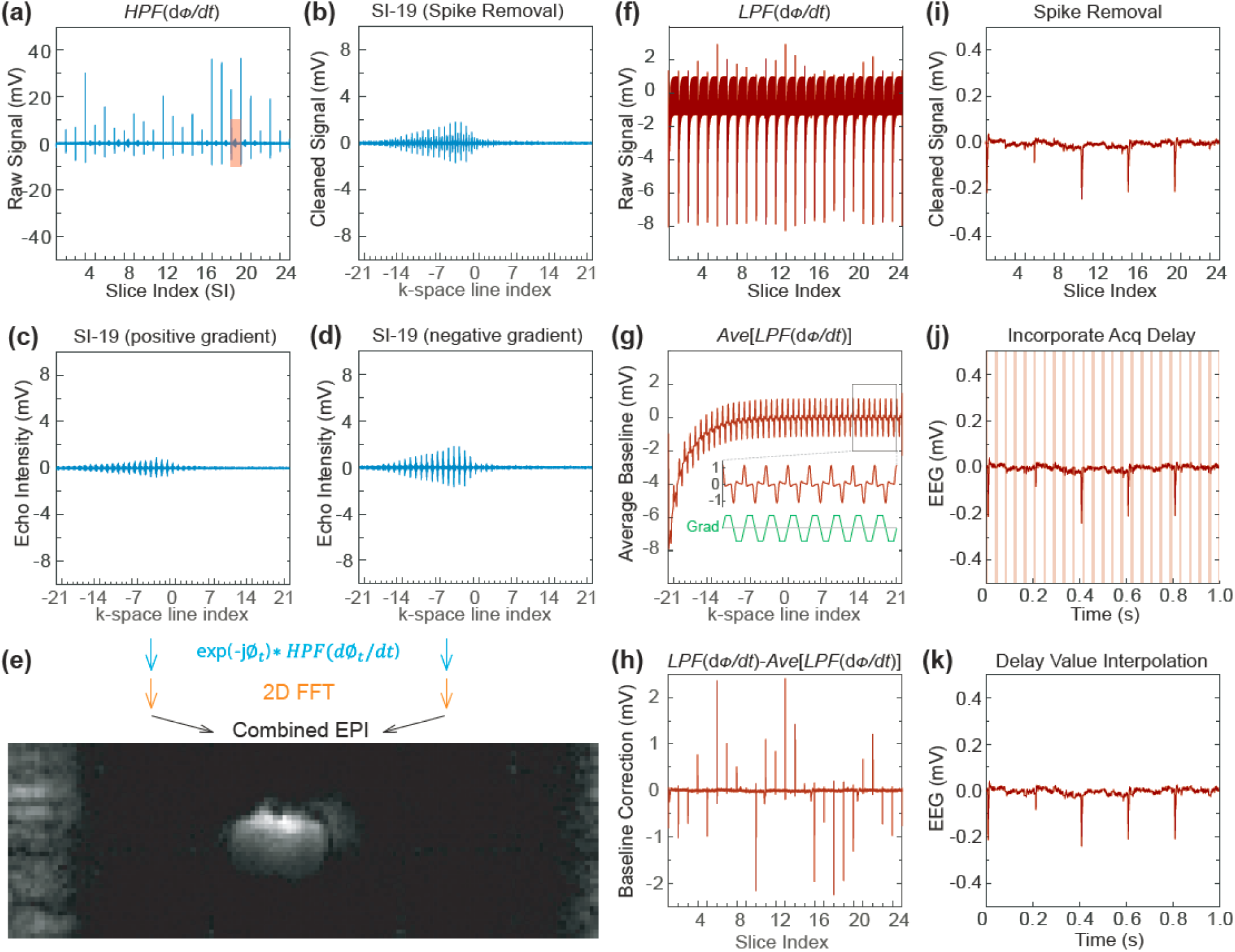
To reconstruct EPI images for all the 24 slices, the detector’s oscillation signal was recorded over the entire duration of the MR acquisition window. The phase ∅_*t*_ of oscillation signal was derivatized over time *d*∅_*t*_/*dt* before being high pass filtered (*HPF*) to obtain the raw signal in (a). The intense spikes appeared between adjacent slices due to acquisition discontinuity. These spikes could be removed by medium filter, leading to cleaned k-space signal, as exemplified by (b) that corresponded to the 19-th slice during interlace acquisition shown by the pink region in (a). To minimize acquisition delays without reducing the effective echo time (TE), each k-space line was sampled during both positive and negative gradients, while the retrieved k-space signals for the positive and negative gradients were separated into (c) and (d). For each point in the k-space, the amplitude signal *HPF*(*d*∅_*t*_/*dt*) was multiplied by the phase term *exp(*-*j*∅_*t*_) of the oscillation signal, leading to a phase-sensitive signal *exp(*-*j*∅_*t*_)∗ *HPF*(*d*∅_*t*_/*dt*) for that time point. This phase multiplication procedure was performed for both signals in (c) and (d) before 2D Fourier transformation was separately performed, leading to 2D images that were combined into a single image frame in (e). To simultaneously retrieve EEG signals, the derivatized phase signal *d*∅_*t*_/*dt* was low pass filtered, leading to raw EEG signal (f) containing repetitive baseline variation pattern that was synchronized with individual slice acquisition. This baseline variation pattern was averaged over all the 24 slices. As shown in (g), the average baseline pattern had 43 pairs of positive and negative peaks, corresponding to 43 k-space lines each of which contained one positive gradient slope and one negative gradient slope. By subtracting each baseline pattern in (f) with the average baseline in (g) multiplied by an empirical factor for optimal cancellation, we obtained the baseline corrected signal (h) that only contained discontinuous spikes at the interface between adjacent acquisition windows. After applying a medium filter to remove spikes, the EEG signal showed up in (i). To convert the horizontal coordinate from slice index in (i) into acquisition time, acquisition delays (pink stripes in Fig. j) were incorporated between adjacent acquisition windows. The EEG value during each acquisition delay was estimated as the average value before and after this delay, leading to continuous EEG pattern. As shown in (k), all optogenetic stimulation pulses started at integral multiples of 0.2s. The negative spike at ∼0.21 s had smaller intensity than other spikes, because part of this spike overlapped with the acquisition delay sandwiched between adjacent acquisition windows for individual slices.

